# Magnetic activation of electrically active cells

**DOI:** 10.1101/2025.02.07.636926

**Authors:** Guillaume Duret, Samantha Coffler, Ben Avant, Wonjune Kim, Angel V. Peterchev, Jacob Robinson

## Abstract

Magnetic control of cell activity has applications ranging from non-invasive neurostimulation to remote activation of cell-based therapies. Unlike other methods of regulating cell activity like heat and light, which are based on known receptors or proteins, no magnetically gated channel has been identified to date. As a result, effective approaches for magnetic control of cell activity are based on strong alternating magnetic fields able to induce electric fields or materials that convert magnetic energy into electrical, thermal, or mechanical energy to stimulate cells. In our investigations of magnetic cell responses, we found that a spiking HEK cell line with no other co-factors responds to a magnetic field that reaches a maximum of 500 mT within 200 ms using a permanent magnet. The response is rare, approximately 1 in 50 cells, but is fast and reproducible, generating an action potential within 200 ms of magnetic field stimulation. The magnetic field stimulation is over 10,000 times slower than the magnetic fields used in transcranial magnetic stimulation (TMS) and the induced electric field is more than an order of magnitude lower than necessary for neuromodulation, suggesting that induced electric currents do not drive the cell response. Instead, our calculation suggests that this response depends on mechanoreception pathways activated by the magnetic torque of TRP-associated lipid rafts. Despite the relatively rare response to magnetic stimulation, when cells form gap junctions, the magnetic stimulation can propagate to nearby cells, causing tissue-level responses. As an example, we co-cultured spiking HEK cells with beta-pancreatic MIN6 cells and found that this co-culture responds to magnetic fields by increasing insulin production. Together, these results point toward a method for the magnetic control of biological activity without the need for a material co-factor such as synthetic nanoparticles. By better understanding this mechanism and enriching for magneto-sensitivity it may be possible to adapt this approach to the rapidly expanding tool kit for wireless cell activity regulation.

## Introduction

Magnetically sensitive cells would enable remote control of cell activity deep inside living organisms. In nature, magnetoreception is believed to be used by animals that need to orient themselves during migrations ^1,2^. However, despite several proposed mechanisms for magnetoreception, none have been effectively recapitulated in genetically modified cells to generate designer magneto-sensitivity. Instead, the targeted magnetic stimulation of excitable cells most reliably resides in the combination of TRP or Piezo channels and nanoparticles that can generate thermal or mechanical stimulation. Synthetic nanoparticles able to generate thermal energy when exposed to RF magnetic fields have been utilized in combination with TRPV1, TRPV4 or TRPA1 to control cells and animal behaviors. In these approaches, exogenic thermo-sensitive proteins are expressed in targeted cells, and synthetic magnetic nanoparticles are delivered to the stimulation site to enable magnetic hyperthermia ^3–5^. The fastest kinetic reported with this magneto-thermal configuration is ∼500 ms for behavioral responses in *Drosophila* under activation of the temperature-rate-sensitive TRPA1 protein ^5^. The stimulation of mechanosensory cells has also been achieved by exploiting the magnetic torques of iron oxide nanodiscs in slowly alternating magnetic fields (5 Hz, 28 mT). This approach allows fast cell stimulation but requires large nanodiscs delivery to the stimulation site (98–226 nm) ^6^.

Alternative approaches have relied on biogenic nanoparticles such as the iron-sequestering protein ferritin combined with TRPV1, TRPV4 or TMEM16 to stimulate cells with static and RF magnetic fields ^7–10^, but the mechanism of action remains unclear ^11,12^. Several explanations have been proposed, including magneto-mechanic ^7,11^, magneto-caloric ^10^, or through the production of intermediate ROS species by ferritin ^8^. The slow changes and prolonged effect on membrane potential over minutes, as opposed to action potentials triggered within milliseconds of stimulation, are compatible with the two latter mechanisms. This long cell response time prevents sub-millisecond modulations, and these constructs have been preferentially applied to slower regulations, such as modulating insulin levels in mice ^9^, impacting embryogenic cell development ^13^, or varying membrane potentials ^8^. Confounding the understanding of magneto-sensitive protein complex mechanisms is the fact that, over the years, studies from various groups were performed using different conditions of magnetic stimulation (field and frequency), temperature, light exposure, and expression systems. The difficulty in adapting this technology to other systems and platforms hindered the translation of these technologies into other labs and the efforts to understand the mechanisms.

Finally, previous experiments have demonstrated that exposing biological samples to static magnetic fields (SMF) for durations ranging from minutes to days can alter neuronal excitatory thresholds and spiking rates,^14–16^, spinal cord conduction ^17^, ion channel dynamics ^18–20^, or gene expression ^21^. These effects have been reported to emerge on the scale of a minute or longer and may extend beyond the exposure period.

Intrigued by the reports of biogenic magnetic sensitivity, we have developed a pipeline to screen the response of thousands of independent cells to an artifact-free stimulation by a permanent magnet through calcium-sensitive imaging and automatic analysis. Using this setup, which produces a magnetic stimulation of ∼500 mT over 200 ms (Fig. 1B), we were surprised to find reproducible fast excitation of spiking HEK cells without adding TRP channels, ferritin, or artificial nanoparticles. We observed responses to magnetic fields in roughly 2% of all cell cultures recorded, but cell cultures that did respond did so reliably to repetitive stimulation. This rapid, sub-second response to a slowly varying magnetic field may share the same underlying mechanism as the longer timescale effects observed upon SMF stimulation in previous studies.

**Figure 1.**
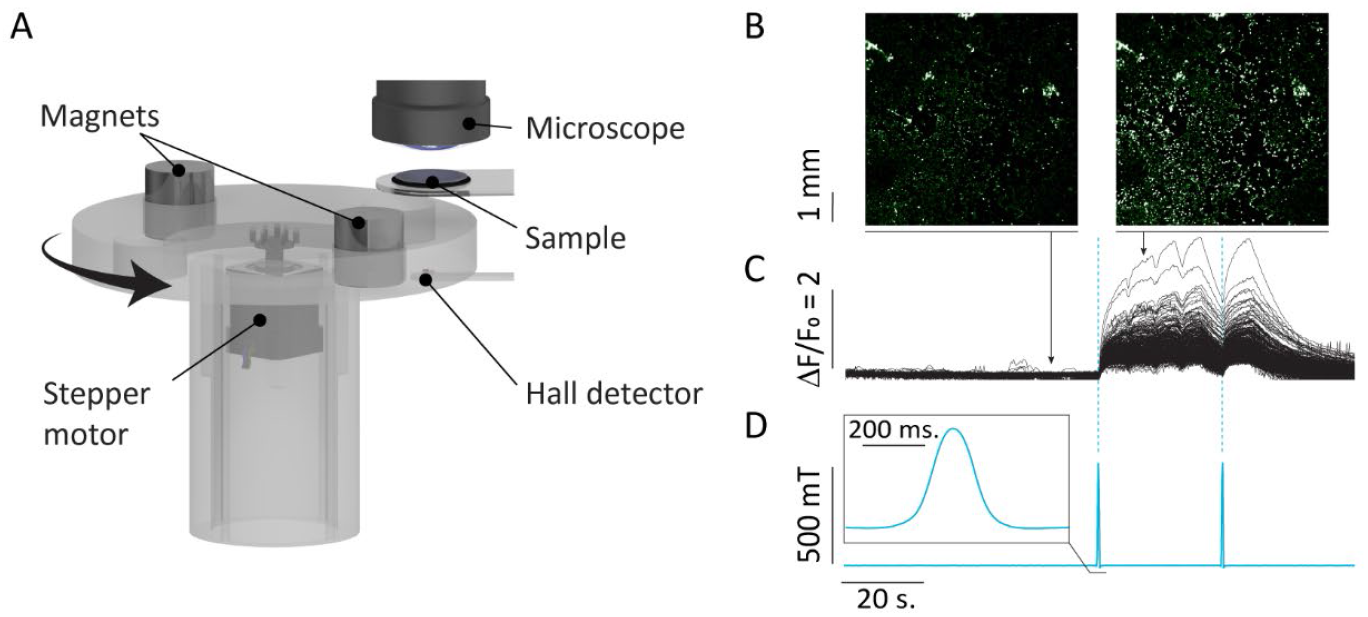
Magnetic-stimulation platform ‘*MagWheel*’. | (A) Our platform consists of a long working distance microscope enabling signal acquisition from over ∼3,000 cells. The stimulation is triggered externally at 60 s and 90 s and monitored with a Hall detector positioned directly below the field of view. The setup is set into a dark enclosure that blocks external light and controls temperature. (B) Calcium fluorescent indicator Fluo-4 is used as a proxy for cell activation. (C) The fluorescent level is computed for each cell using our custom code and plotted as ΔF/F_0_. (D) The applied magnetic field is measured with a Hall detector time aligned to the ΔF/F_0_ traces. The amplitude of the magnetic stimulation is 500 mT and is applied to the sample over less than 200 ms (inset).

## Results

### Magnetic stimulation in a controlled environment

We designed our setup to provide consistent magnetic stimulation under a well-controlled environment that minimizes artifacts during recordings (Fig. 1). Our cell-recording platform is enclosed in a black acrylic box where we can accurately control light exposure and temperature. Using a long working-distance microscope, we can image a 5×5 mm field of view (FOV) and acquire signals from over 3,000 cells. Under the sample, we mounted permanent magnets on an aluminum rotating disc placed on a vibration-dampening platform and controlled the rotating disc using an Arduino-driven stepper motor. We used a Hall-effect sensor to monitor the magnetic field at the sample level during each experiment so we can align the magnetic field intensity with fluorescence imaging. We measured the magnetic fields produced by cylindrical neodymium magnets (DX8×8 N52, K&J Magnets, 38 mm diameter x 38 mm height) to be approximately ∼500 mT at the sample level. During a typical experiment, as the magnets rotate below the sample, the field reaches its maximum value in less than 200 ms, yielding a maximum temporal gradient of 5 T/s (∼10,000 times slower than for TMS), and a spatial gradient of 30 T/m which is orders of magnitude lower than the high-gradient magnetic fields (HGMF; 10^5^–10^7^ T/m) able to apply gravity-like diamagnetic forces to biological tissues (Fig. 1D) ^14,15^. To isolate the magnetic response from artifacts of our experimental apparatus, we designed a sham stimulation consisting of a cast iron block with similar weight and identical dimensions to the neodymium magnet, which we also loaded into the rotating disc. An opaque carbon film is placed between the sample and the magnet to avoid light reflection during stimulation.

Using fluorescent calcium indicators, we could visualize calcium influx into excitable cells, which accompanies action potentials in spiking HEK cells. This is the same approach commonly used to monitor neuronal activity. The fluorescent calcium indicator Fluo-4 requires a 30-minute incubation period before recording but no transfection. This enables the staining of all the cells and a complete activity screening. Moreover, this approach helps extend the protocol to other cells without requiring additional genetic modifications. Furthermore, calcium imaging can dramatically increase the experimental throughput compared to patch clamp electrophysiology, which allows us to detect rare activation events. To ensure that data are fairly compared between stimulation and sham conditions, we developed an algorithm to automatically detect the cells, quantify the fluorescence, and identify peaks in the calcium response (see Methods) ^10^.

### Spiking HEK are sensitive to slowly varying magnetic fields

To study magnetic stimulation of electrical activity, we used modified human embryonic kidney cells (HEKs) that express the voltage-gated sodium and potassium channels NaV1.3 and KiR2.1. These “spiking HEK cells” (spHEKs) provide a well-controlled source of electrically excitable cells, which have been used as a test bed for GEVI development or to study electrical stimulation protocols ^16,17^. Electrical stimulation of spHEKs causes action potentials that propagate to neighboring cells due to the endogenous gap junctions (connexin 43 and 45) that sustain the cells’ electrical couplings ^18,19^. To reduce the background of spontaneous activity that we found at higher temperatures, we performed all recordings at 27 °C. After transferring cells cultured on coverglass to the chamber, we allowed 10 minutes for them to adapt to the chamber conditions before we exposed them to a sham or magnetic stimulation. For each recording, we delivered stimulation at t=60 s and t=90 s for a total calcium imaging acquisition time of 2 min. We waited two minutes between recordings to allow cells to recover between experiments. When we analyzed the calcium response, we found that some cell cultures display a strong calcium influx upon magnetic stimulation that can be repeatedly triggered (Fig. 1B-C). Despite our attention to culture conditions, seeding densities, incubation times, and volumes, we did not observe responses from every cell culture. However, cell cultures that respond to magnetic stimulation once were repeatably activated by additional stimuli, demonstrating the existence of an effective underlying magnetic response in one or many of the cells within the colony. Because the cells are electrically coupled via gap junctions, our single culture recordings cannot reveal whether the calcium activity results from a magnetic response in all or a subpopulation of cells.

We then patterned multiple independent colonies on a single coverslip to increase the number of independent cell cultures monitored during each recording and determine if the majority of cells mediates the response (see M&M). The patterns custom printed on coverslips allow the growth of 400 µm-diameter cell colonies bearing 150-300 cells each, 400 µm apart (side to side), that can be recorded simultaneously (Fig. 2A). In these conditions, when we measured the magnetic responses we found that the majority of cell colonies are *not* activated upon magnetic stimulation, despite being proliferated from the same cell stock. However, as before, colonies that responded to magnetic stimulation did so repeatedly in subsequent magnetic stimulations (Fig. 2B). Importantly, we did not observe any calcium response aligned with sham stimulation, indicating that neither the movement of the wheel nor any artifact from the setup is responsible for these responses.

**Figure 2.**
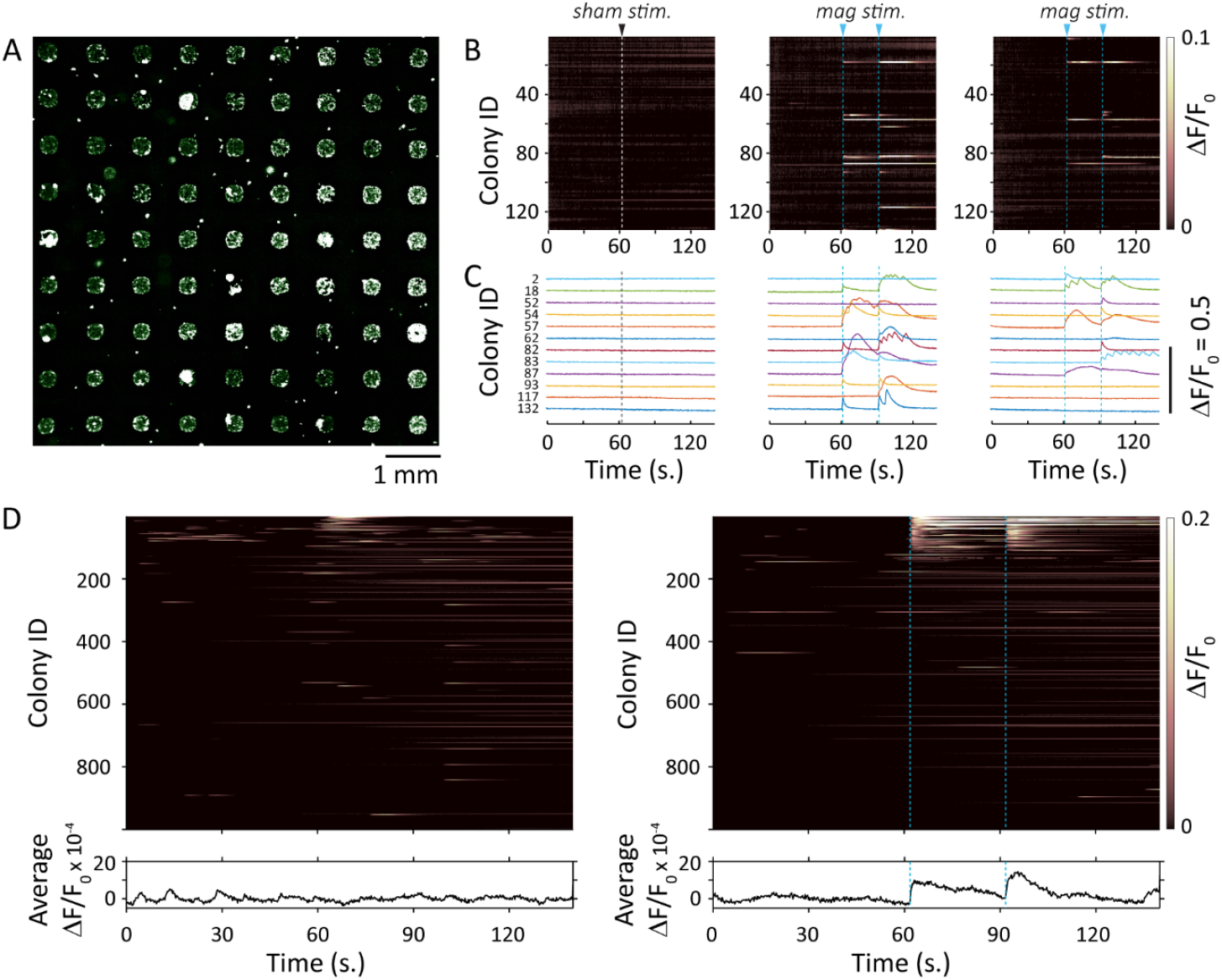
Excitable cells respond to slow magnetic field pulses. A. Spiking HEK cells are plated on a coverslip with printed patterns, allowing the growth of independent 400 µm cell colonies. The cells are loaded with the fluorescent reporter Fluo-4 to record calcium activity. B. Representative recordings computed from 132 colonies of a single coverslip. The calcium activity for each colony is displayed as a heatmap of the fluorescence intensity over time. Each row represents the same colony across all 3 panels. The stimulation time by sham (white line) or 500 mT magnetic field (blue line) is indicated. C. Colonies showing calcium activity within 1 s of magnetic stimulation are identified, and the corresponding fluorescence traces are plotted. Panels A-C show data from a single representative experiment. Panel D gathers the data from 7585 colonies from 66 independent coverslips. For both sham and magnetic stimulation, the data are ordered from strongest to lowest response at 60 s and 90 s, and the highest 1000 responses are displayed.

The frequency of finding magnetically sensitive cells is estimated from our recordings of 66 independent cell cultures on coverslips subdivided into a total of 7585 colonies and representing 1-2 million cells. We calculated the calcium response for each colony by computing the area under the ΔF/F_0_ curve (AUC) for 4 s following each stimulation (60-64 s and 90-94 s). The resulting traces are ranked based on the largest AUC and plotted as a heatmap in Fig. 2D. This heatmap shows ΔF/F_0_ *vs* time for the 1000 colonies with the highest AUCs for sham or magnetic stimulation. Responding colonies are identified by automatically detecting traces displaying a rate change at the time of stimulation. The precise timing of stimulation is confirmed by the hall detector for each recording. Out of the 7585 colonies recorded, 150 colonies responded reliably to the magnetic stimulation, while 29 colonies showed spontaneous activity at the time of stimulation (Fig. S1).

### The response to magnetic stimulation is fast

When spHEKs respond to stimulation by the rotating permanent magnet, calcium influxes start within 200 ms of the stimulation onset. To measure the kinetics of the response, we first select colonies that respond to magnetic fields based on an initial imaging of a large FOV at 5 Hz. Once we identified a responding colony, we increased the magnification (and the frame rate (25 Hz) and repeated the magnetic stimulation. When we plot the inflection point of the calcium fluorescence relative to the magnetic field as recorded with the Hall detector we find that the initiation of the calcium response occurs in the 200 ms window before the peak magnetic field (Fig. 3). This sub-second response to sub-Tesla magnetic stimulation is unlike other magnetic stimulation paradigms we could find in the literature. Previously reported activation mechanisms from similar magnetic gradients and amplitudes require comparably slow mechanisms such as heat transfer or the generation of secondary effectors like reactive oxygen species ^7–9^. For example, during RF magnetic stimulation of cells expressing TRP-ferritin constructs, the ferritin-mediated production of oxygen species and oxidized lipids is ultimately responsible for activating TRP channels and the subsequent change in membrane-proteins dynamics. The response observed in this case occurs seconds after the initial magnetic activation ^8^.

**Figure 3.**
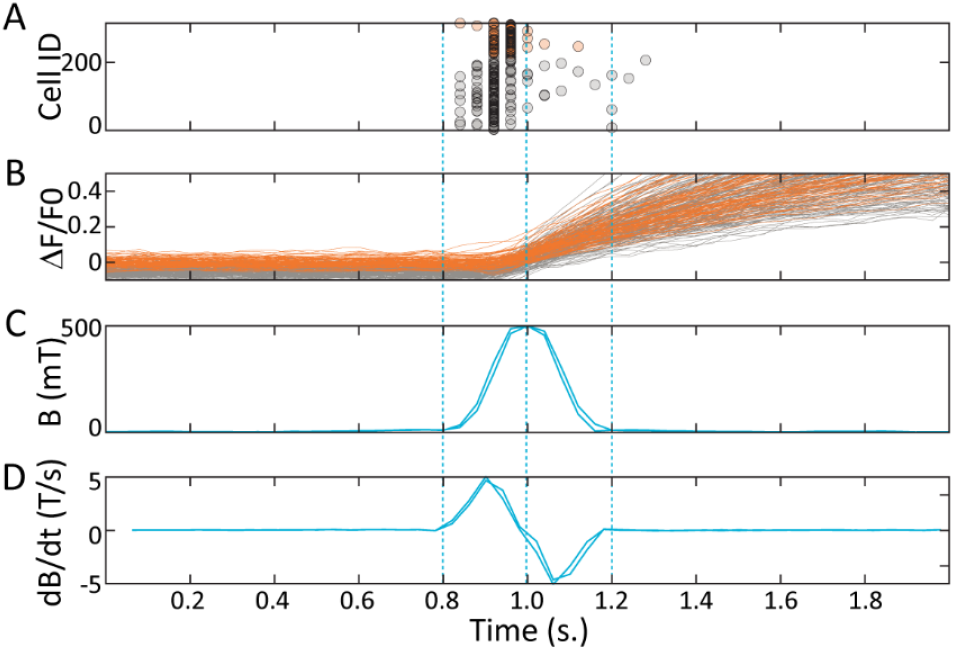
The response to magnetic stimulation is fast. | Responding colonies were recorded at 25 Hz to determine the kinetics parameter of the response to magnetic stimulation. The time of the calcium peak onset is indicated (A) for each trace (B) corresponding to each cell in responding colonies. The corresponding magnetic field strength (C) and corresponding rate of change (D) are measured during stimulation using a Hall detector, and plotted against time. The dotted line indicates the beginning and end of the magnetic stimulus. The calcium peak onset is automatically determined using a custom Matlab script.

### The rapid magnetic response is unlikely to result from the induction of an electric field

Although our stimulation gradient is orders of magnitude slower than TMS (5 T/s *vs*. 20 kT/s), we wondered if this rapid response could be based on electric field induction. Our calculations show that the amount of induced electric field, eddy current, or heat generated at the cell level by this magnetic stimulation is below effective thresholds for either neural activation or modulation of ongoing activity.

Using Faraday’s Law of induction for a quasistatic (low frequency) system ^20,21^, the upper bound on the electric field magnitude (*E*) that can be induced within the cell cultures is

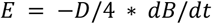

where *D* = 9 mm is the diameter of the recording chamber compartment filled with conductive extracellular buffer solution over the coverslip and *dB/dt* = 5 T/s is the maximum rate of change of the magnetic field over time (Fig. 3C). This results in an estimated maximum electric field induced at the cell cultures of 0.0113 V/m with a biphasic waveform of duration ∼ 0.4 s following *dB/dt* (Fig. 3D). This electric field strength is an order of magnitude lower than the reported minimum to modulate ongoing neural activity (∼ 0.2-0.5 V/m) ^22,23^ and three orders of magnitude lower than the estimated minimum to activate neurons with very long pulses (16.2 V/m) ^24^. The corresponding induced (eddy) current density *(J)* can be estimated at the cell culture level from the induced electric field *E*

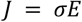

where the measured conductivity is σ= 1.17 S/m. This resultant maximum current density *J* = 0.0132 A/m^2^ is also orders of magnitude lower than used in neural stimulation (tens of A/m^2^) ^24^. Further, the current density can serve to estimate the maximum generated heat (*Q*) and temperature change (*ΔT*) induced by the eddy currents at the cell cultures,

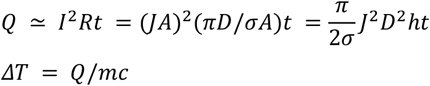

where σ, *h, m, and c* are conductivity (1.17 S/m), height (1.5 mm), mass (9.54×10^−5^ kg), and specific heat conductivity of the extracellular buffer solution (4184 J/kgK), and *t* is time that the field is on (0.4 s), respectively. The resulting *Q* is approximately 1.13×10^−11^ J, and *ΔT* is approximately 2.86×10^−11 o^C per magnetic pulse.

Considering the low calculated values, we ruled out electric field induction and heating as mechanisms, and we decided to explore the implication of another fast-responding mechanism, which is the ability of the cell to respond to mechanical stress through mechano-activated channels such as Piezos and TRPs.

### Mechanosensitivity may explain the magnetic response in spHEKs

Due to the fast kinetic activation, we hypothesize that the magneto-sensitivity in spHEKs may be based on the direct activation of calcium transporters via mechanical effects. Such channels are expressed in HEKs and can be sensitive to transmembrane depolarization (CaV), temperature (nonselective cation TRP channels), or mechanical stress (TRP and Piezo channels). As discussed above, our stimulation is orders of magnitude too weak to impact membrane potential or gate voltage-sensitive channels directly. Therefore, we decided to explore if mechanoresponsive channels relate to magnetic-induced response by selectively inhibiting their mechanosensation.

When one or more colonies on a slide are reliably and reproducibly responsive to the stimulus, we perfuse drugs (or DMSO as a control, Fig. 4A) at the appropriate concentration and record the response to subsequent stimuli. Gadolinium impacts magnetosensitivity by altering the lateral interaction and pressure of lipids in the cell membrane ^25^. At a concentration of 50 µM Gd^3+^, the amplitude of the response to magnetic stimulation is significantly decreased and sometimes eliminated (Fig. 4B). This supports magnetosensation being a mechanosensitive-dependent mechanism in these cells.

**Figure 4.**
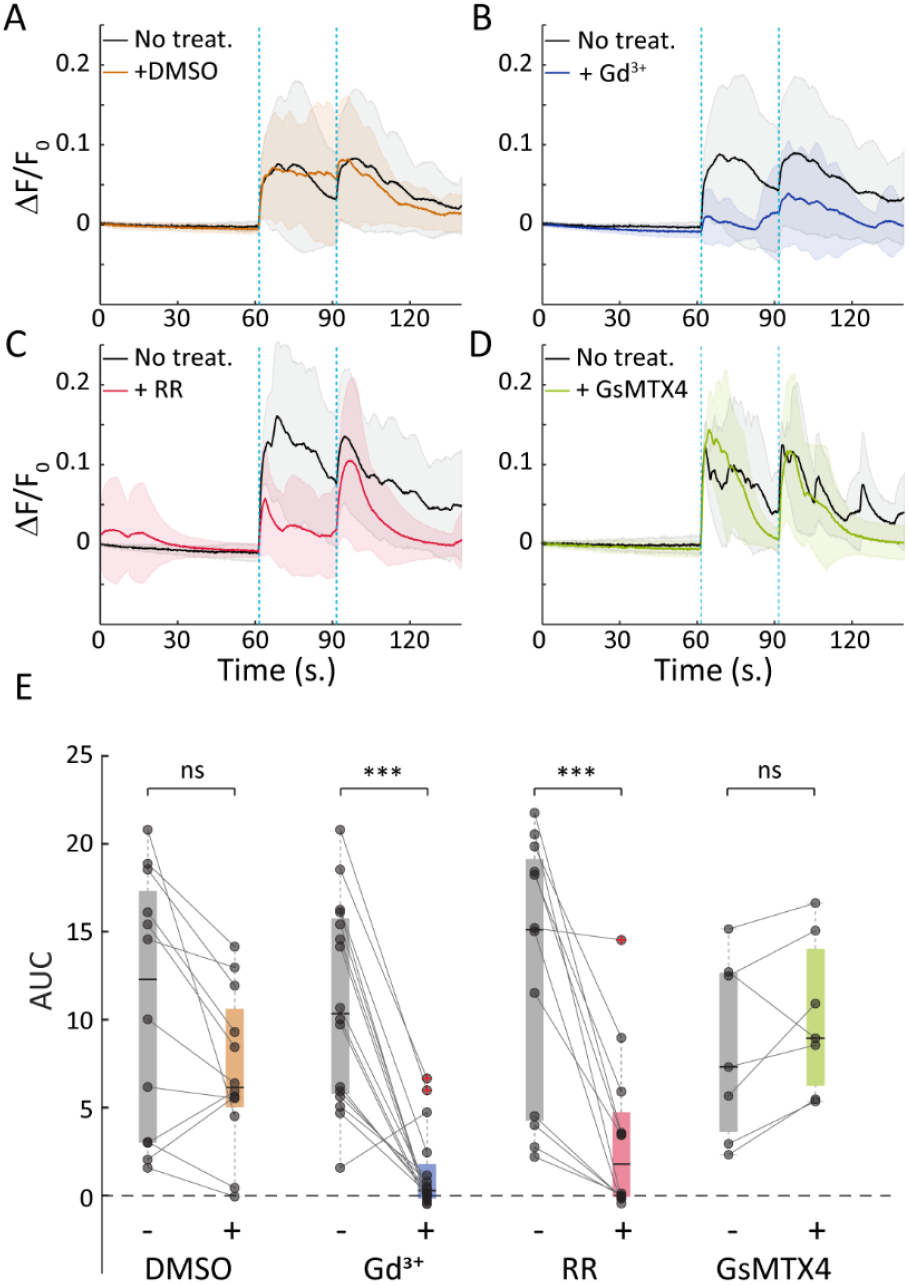
Magnetic response and magnetosensation. | Cells were recorded in the absence (black) or presence (color) of inhibitory drugs for mechanosensitive channels. For each drug, the colonies are recorded first without drugs, and again after adding the drug to the recording bath. Panels **A-D** show the median fluorescent response (solid line) with the box indicating the 25th and 75th percentiles (shaded area), for DMSO (6 slides; 12 colonies), Gadolinium (8 slides; 17 colonies), Ruthenium Red (1 µM: 6 slides 12 colonies), and GsMTX4 (5 slides; 7 colonies). Panel **E** summarizes the data by computing the area under the curve (AUC) after magnetic stimulation for each responding colony, before (−) and after (+) exposure to the indicated inhibitors or to the DMSO control. (Anova; ns: P ≥ 0.05; ^*^ : P < 0.05; ^**\*\***^ : P < 0.01; ^**\*\*\***^ : P < 0.001).

**Figure 5.**
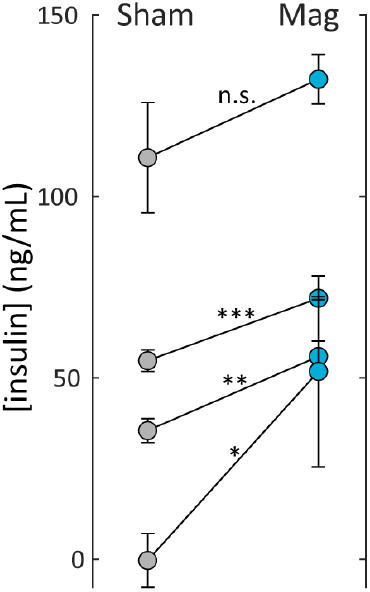
Activation of insulin secretion by magnetic stimulation. | spHEK cells and MIN6 cells are grown together and exposed to sham or magnetic stimulation. After 1 hour, the media is collected and insulin is quantified by ELISA. Each data point shows the average content (+/-sd) from 3 wells seeded and stimulated simultaneously on the same 24-well plate. (paired t-test; **ns**: P ≥ 0.05; ^**\***^ : P < 0.05;^**\*\***^ : P < 0.01; ^**\*\*\***^ : P < 0.001).

We decided to investigate further, and specifically inhibit the major proteins involved in sensing mechanical changes at the membrane level. Most TRP channels are mechanosensitive, and all are permeable to cations ^26^. The molecule Ruthenium Red (RR) can block channels of the TRPV subfamilies, including the mechanosensitive TRPV1, TRPV2, or TRPV4, which are expressed in HEK cells (IC50 ∼500 nM) ^27,28^. The perfusion of a 1 µM RR solution significantly decreases the response amplitude compared to pre-perfusion recordings, suggesting the involvement of these channels in the translational mechanism permitting magnetosensation in spHEKs (Fig. 4C and E). The partial inhibition of the response could be due to the additional implication of TRP-independent mechanisms or to the activation of TRP channel at the level of the calcium-loaded endoplasmic reticulum (ER). We then investigate the impact of the Piezo inhibitor GsMTX4 (20 µM). The pre- and post-incubation recordings show similar response amplitudes, which suggest that Piezo channels play no role in the mechanism of action.

Here, we demonstrate the implication of the mechanosensitive channels of the TRP family in the response mechanism. At this point, we cannot reliably determine whether the TRP channels are intermediate or primary responders. A response below 200 ms after the magnetic stimuli onset supports a direct mechanism devoid of time-consuming secondary actors ^29^.

### Magnetic torque on lipid rafts could explain the magnetic response

Structures with regular arrangements, such as cell membranes and cytoskeleton proteins, can have diamagnetic anisotropy in magnetic fields ^30^. Magnetic fields can force these cellular elements to align with the field direction and exert torque on associated or embedded proteins and channels. This can particularly affect the orientation and tension of the cell membrane and increase the mechano-sensitive channels’ open probability ^15,31^. The magnetically induced mechanical forces at the cell level can be estimated based on the principles of magneto-mechanical interactions ^11,31^. The volume density of the magnetic gradient force (*f*) acting on a cell is given by:

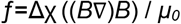

where *Δχ* is the difference between the susceptibility of the medium and cells (−1.8 ·10^−6^) ^32^, *B* is the magnetic flux density, and *μ*_*0*_ is the permeability of free space (4π×10^−7^ H/m). With a magnetic field strength of 0.5 T and a spatial gradient of 30 T/m, the magnetic gradient force density *f* is ∼22 N/m^3^, which results in a force of ∼2.7×10^−20^ N at the cell membrane level. To approach the pN range of force needed to gate mechanosensitive channels directly ^33^, we would need a gradient of 10^8^ T/m for a 1 T magnetic field.

However, discreet effects on the membrane surrounding TRP channels might have a strong impact. Lipid rafts are cholesterol-rich microdomains that play a significant part in mechanosensation through their interaction with TRP channels ^34^. These liquid-ordered domains have larger dipole potential ^35^ and different magnetic susceptibility than the surrounding lipid bilayer ^15,36^. Their alignment along the magnetic field in regard to the rest of the membrane can alter the activation and inactivation kinetics of associated TRP channels, notably through modification of the lipid bilayer curvature ^37,38^. One way to test this hypothesis would be to modify the lipid membrane’s dielectric properties and lipid rafts’ occurrence to modulate the spHEK cells’ magneto-sensitivity.

### Magnetic stimulation of spHEK co-cultures drives insulin production

Although magnetic stimulation of action potentials in spHEKs is rare, we hypothesized that an activated cell could catalyze the activation of a network of cells electrically coupled via gap junctions. In this way, the small number of magnetically sensitive cells could induce calcium-dependent activities in a heterogeneous cell assembly. We tested the ability of spHEKs to act as sensitizers to magnetic fields by co-culturing them with beta-pancreatic cells MIN6.

In the presence of glucose, MIN6 cells release insulin to the extracellular environment when subjected to depolarization or calcium influx. We co-plated 40,000 spHEK cells and 120,000 MIN6 cells on coverslips and let them grow together for 72 hours. Following a 30-minute incubation in starving media (DMEM with 1 mM glucose), we exposed the co-culture to 3 sessions of 10 magnetic stimulations (0.5 Hz) or sham stimulation in the presence of 3 mM glucose. Following stimulation, we incubated the cells for 1 hour at 37 °C, and collected the media. When we plotted the insulin concentration, measured by ELISA, for cells exposed to sham and magnetic stimulation, we found that in all cases, the magnetic stimulation increased the insulin levels relative to sham stimulation. These results demonstrate that despite an estimated response of only about 2 % of spHEKs to magnetic stimulation, that level of magnetic sensitivity is strong enough to modulate the activity of an associated cell population.

## Discussion

We report the fast activation of excitable cells by a time-varying non-uniform magnetic field, yielding a maximum magnetic flux density of 500 mT, with a spatial gradient of 30 T/m and a temporal gradient of 5 T/s. To our knowledge, such stimulus has never been reported to induce nearly immediate (< 200 ms) action potentials in electrically excitable cells without artificial or biogenic nanoparticles.

### Similarities to previous studies and other potential mechanisms

Decades of research have studied the impact of SMF on cells and biological tissues. For example, applying a strong permanent magnet to the human cortex induces neurophysiological and behavioral effects, and could lead to clinical applications for transcranial static magnetic field stimulation (tSMS) ^39^. However, effects have mainly been reported as relatively small changes in cell excitability ^12,40,41 42^, with effects persisting for seconds to minutes after exposure ^14,15,31,43–50^. SMF has also been shown to impact cell and tumor growth, ROS levels, cytoskeleton, and cell migration ^42^, with a kinetic that correlates with magnetic spatial gradient rather than flux density. Gradients of 10^3^ T/m can trigger apoptosis over days ^51^, while a 10^6^ T/m gradient can effectively remodel the cytoskeleton in hours ^32^. And if high magnetic gradient stimulation could reach 10^9^ T/m, we expect the resting membrane potential to be impacted in milliseconds ^31^.

The effects of SMF can be attributed to the reorientation of biomolecules, such as microtubules, actin filaments, or lipids, along the applied magnetic fields ^52,53^. The diamagnetic anisotropy of lipid bilayers can explain, for example, the silencing of the bacterial mechanosensitive channel MscL seconds after constant exposure to a ∼400 mT SMF ^54,55^. This illustrates how cell membranes can translate even modest magnetic stimuli into mechanical stimuli, significantly impacting cell functions over long periods. This membrane-dependent mechanism has been proposed to explain many other SMF effects ^48,53,56^, including the decrease in MEPP frequency in synapses or the increase in the activation time constant of voltage-activated sodium channels recorded seconds (20-50 s) after the 100 mT magnetic field onset ^57,58^.

Our experiments may be consistent with previous SMF findings in that we hypothesize the response results from the movements of membrane clusters such as lipid rafts. However, unlike previous SMF studies that reported subtle changes in excitability, we observed a much faster response and the propagation of induced action potentials.

Change in the concentration of free radicals is another mechanism by which magnetic fields can affect cell activity, but this mechanism is much slower than the responses we observe ^59^. Evidence suggests that changes in ROS concentration can explain the magnetic sensitivity of cells expressing artificial TRP-ferritin constructs when exposed to time-varying, nonhomogeneous magnetic fields ^60,61^. However, this TRP-ferritin approach for magnetosensitivity results in membrane potential alterations seconds after the stimulation. This response delay results from the dependence of this pathway on magnetically induced secondary messengers, notably ROS ^8,60^. Although these constructs have been employed successfully for neuro-modulation in vitro and in vivo ^8,60,62^, the response latency is several seconds and unlikely to explain the sub-seconds action potential generation we report here.

Finally, magnetic-dependent fluorescence is another mechanism that can modify molecular behavior on the millisecond time scale. For example, the fluorescence brightness of EGFP can be modulated within milliseconds of exposure to a 25 mT magnetic field. This rapid and reversible change is postulated to result from modulation of the electron transfer rates in the excited and ground state in the presence of a magnetic field. This hypothesis is sustained by the addition of flavins, which can accept and donate electrons, enhancing this magnetic sensitivity ^63^. This mechanism is unlikely to explain channel gating and calcium influx, as charge transfer within the molecule is not expected to affect open probability.

To our knowledge, the only other direct magnetic cell stimulation without magnetic materials that shows response times in the millisecond range is transcranial magnetic stimulation (TMS). TMS can generate action potentials within 1 ms after stimulation, as it acts through induced electrical stimulation ^64–67^. However, this method relies on a high temporal gradient - 20,000 T/s compared to 5 T/s for our setup - able to induce electric fields and depolarize the targeted cells. TMS-like mechanisms also cannot explain why only some cells consistently respond to magnetic stimuli when all cells are experiencing similar electric fields. We report here that some cells can respond to a sub-tesla slowly varying magnetic field while other neighboring cells remain silent. This caveat, although impairing a better efficiency for now, will constitute a clear advantage if we can purposefully make specific cells more sensitive than others through genetic manipulations. We can then modify neurons in vivo to make them receptive to the magnetic stimulus or genetically engineer cells for injectable cell-based therapeutics.

### Applications and future work

Although more work is needed to validate the mechanism of action, we can exploit this magnetic sensitivity to regulate cell activity. This utility is illustrated by the co-culture experiment where the activation of the spHEK cells by magnetic stimulation depolarizes the nearby pancreatic cells, leading to a calcium influx and insulin secretion. In this case, increasing the number of magnetic stimulations (3 × 10 pulses) and using a denser culture allowed any responding (spHEK) cell to stimulate nearby (MIN6) cells, resulting in insulin production. One route for future work is to adapt this concept for in vivo experiments and therapies through injectable encapsulated co-cultures that we can control remotely with a magnetic field, similar to prior work that relied on magnetic nanoparticle cofactors ^68,69^.

Another important research direction is to increase the number of responding cells, which a complete understanding of the underlying mechanism will aid. Based on our hypothesis involving lipid rafts, it may be possible to modulate the density of those microdomains and, thus, the likelihood of magneto-sensitivity in spHEKs. Increasing the number or sensitivity of mechanoreceptors in the cell membrane may also increase the response probability of spHEK to magnetic fields. We have attempted to overexpress TRP channels but without success, potentially due to these manipulations affecting cell health. It may be possible to modify the expression of TRP or other mechano-sensitive channels, such as eMscL or Piezo, in ways that maintain cell health.

Overall, our data shows that sub-tesla magnetic fields delivered in roughly 200 ms can generate action potentials in cells without additional magnetic co-factors. While the mechanism is not completely understood, evidence suggests it follows a mechanical stimulation pathway. By better understanding the mechanism and enhancement of the cellular response, it may be possible to control cell activity remotely with magnetic fields that can be generated with a permanent magnet.

## Materials and Methods

### Cell culture

Spiking HEK293 cells (gift from Adam Cohen’s laboratory) are grown in DMEM-F12 (Lonza) supplemented with 10% FBS (Gibco) and 1% penicillin-streptomycin (Lonza) at 37 °C in 5% CO_2_ atmosphere conditions. The cells are passaged at <70% confluence for 10 generations or less before being replaced with a fresh stock. We observed a decrease in spontaneous and induced activity in cells with a higher passage number or cells that have been grown to a higher confluency. For stimulation and recordings, cells are replated on 12 mm glass coverslips at 26,000 cells / cm^2^ (50,000 cells / well). The coverslips are previously flamed and when needed, patterned. The response to magnetic stimulation is recorded 60-72 hours after replating.

MIN 6 cells (ATTC) are grown at 37 °C (5% CO2) in DMEM-F12 (Lonza) supplemented with 15% FBS (Gibco), 55 µM β-mercaptoethanol and 1% penicillin–streptomycin (Lonza). These cells are passaged every 48 hours at a 1:2 ratio. For the co-culture experiments, the 15 mm wells of a 24 well-plate are seeded with 40,000 spHEK cells and 120,000 MIN6 cells. After 72 hours, the cells are starved for 30 minutes in DMEM-F12 containing 1 mM glucose. The media is then replaced with DMEM-F12 containing 3 mM glucose, and the cells are transported to the MagWheel platform for stimulation. The cells are brought back to the incubator for 1 hour, and the media is collected for insulin quantification.

### Fluorescent calcium imaging and magnetic stimulation

All experiments are performed in extracellular buffer (ECB: 119 mM NaCl, 5 mM KCl, 10 mM HEPES, 2 mM CaCl_2_, 1 mM MgCl_2_ (pH 7.2); 320 mOsm). To record calcium influx, the cells are incubated in culture media with 2 µM Fluo-4-AM (ThermoFisher) for 30 minutes. The coverslip with the cells is rinsed with ECB and transferred to the recording chamber with 100 uL of ECB. The aperture of the chamber forms a 9 mm diameter opening above the coverslip, resulting in a 1.5 mm height of the ECB above the coverslip and the cells. The cells are left to equilibrate for 10 minutes at 27 °C in the microscope enclosure prior to imaging. An AirTherm™ SMT (WPI) fitted to the microscope enclosure maintains and controls the temperature to within 0.2°C. The recording chambers are 3D printed to mimic Warner QR-48LP models but without the magnetic and metallic elements.

Imaging is performed with a Nikon SMZ18 epifluorescence stereo microscope. We use a long working distance 2X objective (SHR Plan Apo; wd 20) to allow imaging of a large field of view (5×5 mm) and prevent interactions with the magnetic field. Images are collected with a Zyla sCMOS Camera (Andor, Belfast, UK) through a GFP Filter Cube Set (Nikon). The acquisition is set for 700 frames at 5 frames per second (fps) (200 ms exposition, 140 s total). To lower the file size and increase the signal, we bin 2×2 pixels to obtain a 1020×1020 pixel image. Image acquisition triggers the magnetic wheel to start 30 s later and coordinates the hall detector acquisition rate to the frame rate. At the end of the recording, the file is saved as a 16-bit TIF file, with no compression or modification. For the kinetic analysis, 1000 frames are acquired at 25 fps, and the stimulation is triggered at t = 10 s.

The magnetic stimulation is delivered with a N52 1.5 in x 1.5 in (38 mm x 38 mm) DX8×8-N52 Neodymium Cylinder Magnet (K&J Magnetics) mounted on an aluminum disc. The disc is rotated on demand using a stepper motor controlled with an Arduino. The Arduino is also connected to a Hall detector mounted directly under the sample to monitor the magnetic field at the sample level for every recorded frame. This also provides a feedback loop to stop the stepper motor after a single magnetic stimulation. For each recording, the stimulation is delivered at t=60 s and t=90 s for a total acquisition time of 2 min. To allow cells to recover, full recordings are set 2 min apart. The whole setup is enclosed in a black acrylic box, where the temperature is controlled, and light cannot penetrate.

### Patterning coverslips

Hydrogel printing is based on the 3D stereolithography techniques previously published ^70^. The pre-hydrogel solution contains 20% wt 3.4 kDa poly(ethylene glycol) diacrylate (PEGDA), 34 mM lithium phenyl-2,4,6-trimethylbenzoylphosphatine (LAP) as the photoinitiator, and 2.25 mM tartrazine as the photoabsorber in PBS. 100 uL of the pre-hydrogel solution is placed on top of a flamed coverslip. Using a Lumen 3D bioprinter, targeted polymerization is catalyzed using 405 nm light projected through a digital photomask. The resulting pattern consists of 400 µm-diameter wells separated by 400 µm walls of crosslinked PEG-DA. Following irradiation, the coverslips with hydrogels are rinsed in PBS 2 times, sterilized with UV,, and stored in PBS at 4 °C until use (< 3 weeks).

### Computing onset of peaks, AUCs, DF/F0

The collected fluorescence images are analyzed using custom algorithms developed in MATLAB (The MathWorks, Natick, MA). Regions of interest (ROIs) corresponding to individual transfected cells or to individual colonies are automatically selected through a segmentation algorithm. For each ROI, the change in fluorescence (ΔF/F_0_) is calculated based on the average fluorescence value divided by the average fluorescence value of the first captured image, F_0_. ROIs are classified as responsive if a rate of change in fluorescence is detected within a time window spanning from 400 ms before the peak of the magnetic field (measured with the Hall detector for each recording) to 1 s after. This is done by applying a peak detection algorithm to the derivative of the ΔF/F_0_ traces acquired with magnetic and sham stimulation (see Fig. S1). The algorithms used in this publication can be shared, please contact the authors.

## Supplementary Material

**Figure S1.**
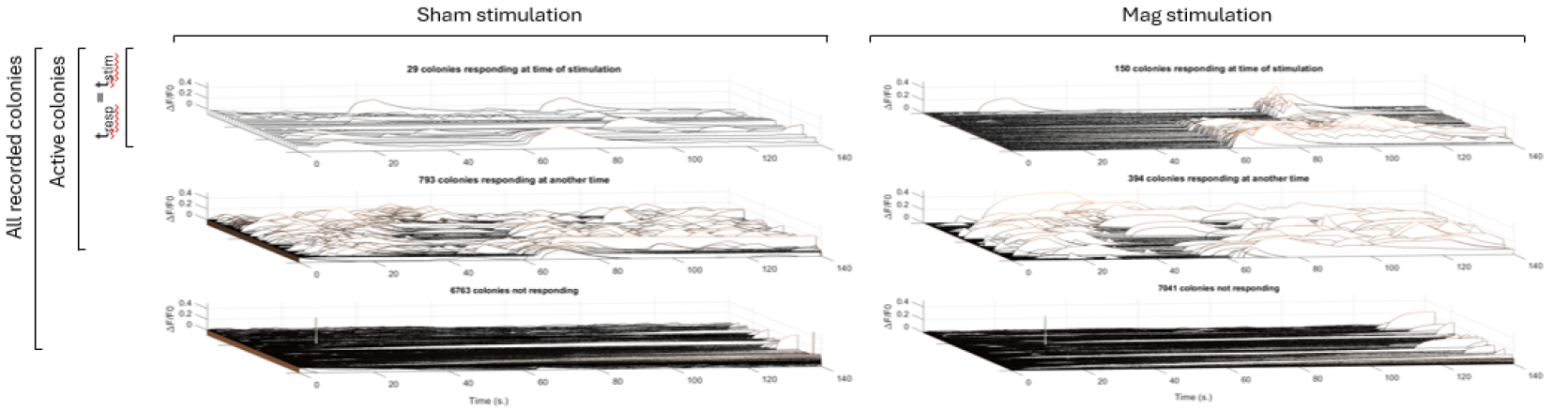
Calcium activity traces for each colony. | The ΔF/F_0_ traces recorded for each colony are derived against time, and peak detection is utilized to detect rate changes (*t*_*resp*_). The time of stimulation *t*_*stim*_ is a time window from 400 ms before the peak of the magnetic field (measured with the Hall detector for each recording) to 1 s after. For Sham stimulation, the average timing of magnetic stimulation (61.4 s and 91.4 s) is chosen to establish the analysis time windows ([61-62.4] s and [91-92.4] s). For a total of 7585 colonies recorded, 29 colonies exposed to the sham stimulation have some calcium activity (response time *t*_*resp*_) at the time of stimulation *t*_*stim*_. This number is 150 colonies for magnetic stimulation. The rest of the colonies either show activity outside of the time of stimulation (793 and 394 colonies for sham and magnetic stimulation, respectively) or no detected activity (6793 and 7041 colonies for sham and magnetic stimulation, respectively).

## Acknowledgments

The authors thank Joseph Asfouri, Gabriella Franco, Jeanette Ingabire, Kelly Kim, and Charles Sebesta for useful discussions on the effect of magnetic fields. The authors thank Caroline Griffin for all her administrative support and the Rice University Machine Shop for help with manufacturing. This research was developed with funding from the National Institutes of Health NIH grant 1RF1NS126063 / 180014, and the Defense Advanced Research Projects Agency (DARPA), contract N6600119C4020 / R1A260-A.

